# Differences in non-linearities determine retinal cell types

**DOI:** 10.1101/2022.05.26.493557

**Authors:** Francesco Trapani, Giulia Spampinato, Pierre Yger, Olivier Marre

## Abstract

Classifying neurons in different types is still an open challenge. In the retina, recent works have taken advantage of the ability to record a large number of cells to classify ganglion cells into different types based on functional information. While the first attempts in this direction used the receptive field properties of each cell to classify them, more recent approaches have proposed to cluster ganglion cells directly based on their response to standard stimuli. These two approaches have not been compared directly. Here we recorded the responses of a large number of ganglion cells and compared two methods for classifying them into types, one based on the receptive field properties, and the other one using their responses to standard stimuli. We show that the stimulus-based approach allows separating more types than the receptive field-based method, leading to a better classification. This better granularity is due to the fact that the stimulus-based method takes into account not only the linear part of ganglion cell function, but also non-linearities. A careful characterization of non-linear processing is thus key to allow functional classification of sensory neurons.

## 1 Introduction

A striking feature of neural systems is their diversity. To make sense of this diversity, a necessary step is to have a “parts list”, a taxonomy of the different cell types in the brain. To do so, one must cluster neurons in several types, i.e. groups with uniform properties. This is still an open challenge for many parts of the brain [20, 15, 12, 25]. In the retina, several studies have tried to determine what are the different types of retinal ganglion cells (RGCs), the retinal output. Retinal ganglion cells can be classified according to their genetics, anatomy, or physiology. Each type is supposed to extract a specific feature from the visual scene. The most recent studies have shown that there are probably 30 to 40 different types of ganglion cells in the mouse retina [22, 19, 1, 23].

A commonly used approach to define these cell types relies on their function, i.e. on the characterization of how ganglion cells respond to stimuli. Such an approach is especially relevant when the driving purpose is to understand what is the specific computation carried out by each type [23]. The first studies that tried to define ganglion cell types based on their physiology relied on receptive fields: cells with similar receptive field properties were clustered together and considered as part of the same functional groups [18, 21, 6, 17]. More recent attempts aiming at performing large-scale classifications of ganglion cell types relied directly on their responses to a variety of standard stimuli [1, 3, 5, 9]. A key hypothesis in these more recent methods is that two neurons of the same type should extract the same feature, but in a different location of the visual space. For this reason, they should respond the same way if a stimulus that is spatially uniform is displayed [1], or if the stimulus is re-centered on the receptive field for each cell [3, 5, 9, 11]. One of the most recent attempts to classify all ganglion cell types based on their function used, among other features, the responses to this spatially uniform “chirp” stimulus to divide cells in different types [1]. These two classes of methods, the one based on receptive field properties, and the one based on direct responses, have not been compared directly. One would assume that using responses to dynamical stimuli allows to extract more interesting features than just measuring receptive fields, and should therefore allow a more precise classification. However, this has never been tested.

Here we compared two methods for cell type classification in the retina and grouped the retinal ganglion cells recorded in mice retina. The first method, called “RF-based” method, used only the ganglion cell receptive fields to group them in types. The second one, hereafter called “chirp-based” method, also used their response to the so-called “chirp” stimulus.

In a first part, we show that the chirp-based method manages to divide the population into a larger number of cell types, and thus find some types that would have been missed (or not distinguished) by the RF-based method. Then, we asked why this chirp-based method was able to identify more types. We found that the chirp stimulus evoked nonlinear responses in ganglion cells, and that this nonlinear component was necessary to single out some specific types. Our study suggests that a functional classification of neurons require characterizing their non-linear behaviours.

## 2 Results

### 2.1 Comparison of two methods for functional classification of RGCs

Our first purpose was to use the “chirp-based” approach to cluster a large ensemble of ganglion cells into several groups corresponding to putative types. We recorded the spiking activity of ∼500 ganglion cells from 6 different experiments performed on 6 different animals (see Methods). While recording, we displayed a checkerboard to map the receptive fields, and a chirp stimulus with varying temporal frequencies and contrasts (fig. 1.A). This chirp stimulus has already been used previously in several studies to divide populations of ganglion cells in different cell types based on their responses [1, 8]. However, in these studies, ganglion cell activity was recorded with 2 photon imaging, while in the present study, we recorded the activity of the ganglion cells with extracellular recordings and high density Multi-Electrode Arrays (MEAs) (see Methods).

**Figure 1:**
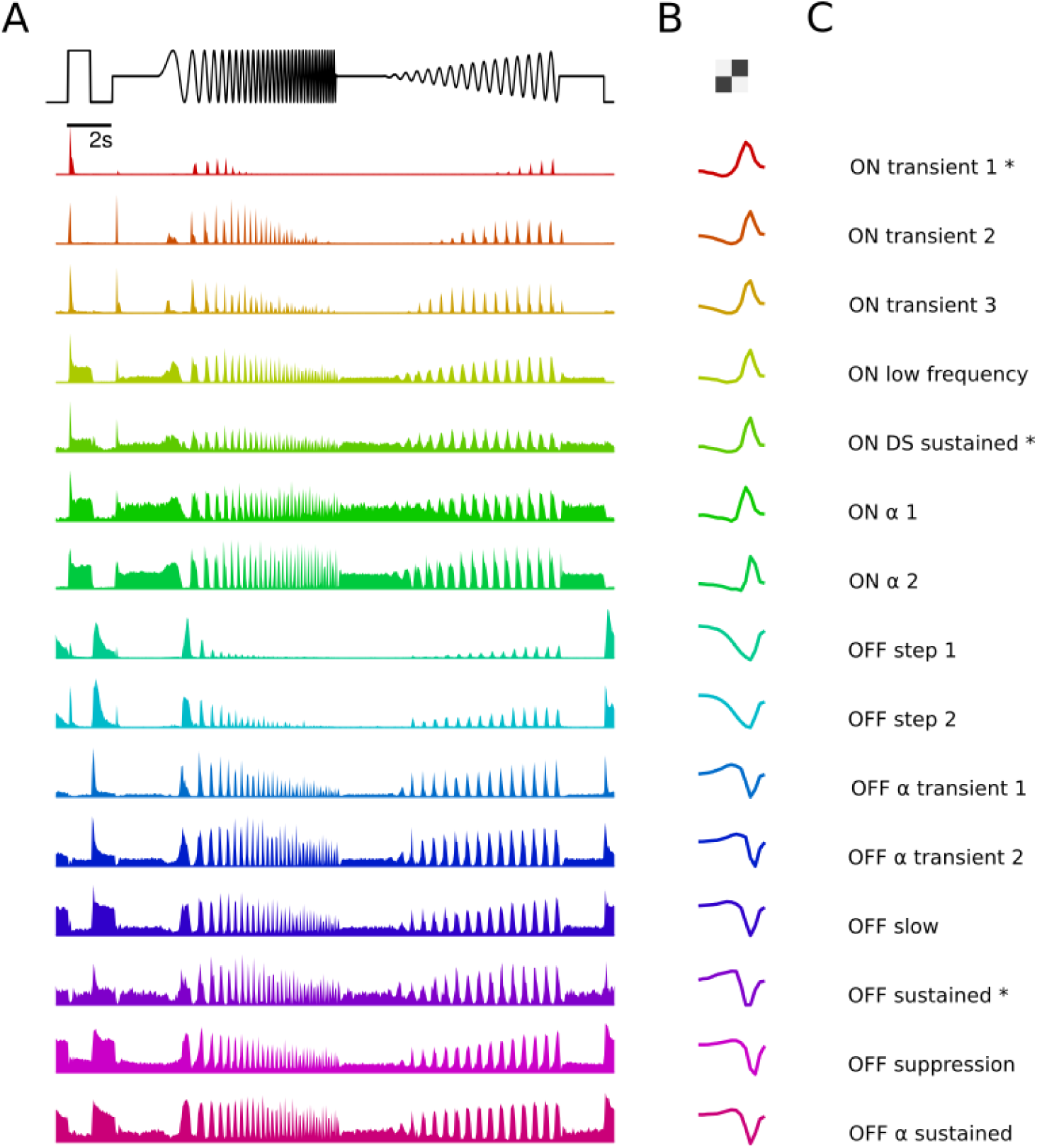
Results of the chirp-based method. The 15 RGC clusters identified and validated with the chirp-based method. **A**. On top, the temporal profile of the chirp stimulus. Below, for each cluster, the cluster-mean response to the chirp stimulus. **B**. For each cluster, the cluster-mean temporal component of the receptive field. **C**. The biological type corresponding to each cluster. Asterisks indicate clusters for which the type is uncertain.

We first selected ganglion cells that had a detectable receptive field, and responded reliably to the chirp stimulus. We characterized each cell with its response to the chirp, the diameter of its receptive field, and the temporal profile of its receptive field. This corresponds to a vector with 520 coordinates for each ganglion cell (see Methods). We first projected the ensemble of vectors onto a subspace of reduced dimension with principal component analysis (PCA, see Methods), and then clustered it into different groups following a method similar to [1]. We obtained 24 clusters, where responses and receptive fields inside each cluster were homogeneous (fig. 1.A, B, C, see Methods). To test if these clusters corresponded to real types, we relied on an established validation criterion [21, 17]: if a cluster corresponds to a single type, the receptive fields of its ganglion cells should form a mosaic tiling the visual space. For each cluster, we grouped together cells belonging to the same retina, and quantified their tiling by measuring the normalized distance between each pair of nearest neighbours. We estimated the distribution of these distances for each group, and tested if there was a significant peak entailing uniform spacing (fig. 2.A, B, see Methods). We found 15 valid mosaics, each corresponding to a different cluster, and all composed of 5 or more cells (table 1).

**Table 1:** Results of the chirp-based method. Summary of the types found with the chirp-based method and validated with the mosaic test. For each type we report: its name (RGC type), the size of its cluster, the number of cells composing its mosaic, its recording bias, the corresponding type from the [1] taxonomy ([*Ca*^2+^] match), and its similarity to the [*Ca*^2+^] match (*ρ*[*Ca*^2+^])

| RGC type | cluster size | mosaic size | recording bias ($\Phi$) | $[Ca^{2+}]$ match | $\rho [Ca^{2+}]$ |
| --- | --- | --- | --- | --- | --- |
| ON trans. 1 | 10 | 8 | 1.46 | N/A | N/A |
| ON trans. 2 | 12 | 10 | 1.63 | ON trans. | 0.82 |
| ON trans. 3 | 14 | 13 | 0.88 | ON trans. | 0.81 |
| ON low freq. | 16 | 10 | 2.23 | ON low freq. | 0.73 |
| ON DS sust. | 9 | 5 | 1.17 | ON DS sust. 3 | 0.65 |
| ON $\alpha$ 1 | 8 | 6 | 2.69 | ON $\alpha$ | 0.73 |
| ON $\alpha$ 2 | 16 | 9 | 2.29 | ON $\alpha$ | 0.69 |
| OFF step 1 | 26 | 7 | 2.01 | OFF step | 0.64 |
| OFF step 2 | 31 | 6 | 1.26 | OFF step | 0.58 |
| OFF $\alpha$ trans. 1 | 27 | 12 | 1.85 | OFF $\alpha$ trans. | 0.80 |
| OFF $\alpha$ trans. 2 | 10 | 5 | 0.78 | OFF $\alpha$ trans. | 0.79 |
| OFF slow | 18 | 7 | 1.90 | OFF slow | 0.62 |
| OFF sust. | 9 | 9 | 2.62 | N/A | N/A |
| OFF suppr. | 15 | 12 | 0.68 | OFF suppr. 2 | 0.76 |
| OFF $\alpha$ sust. | 7 | 5 | 1.73 | OFF $\alpha$ sust. | 0.74 |

**Figure 2:**
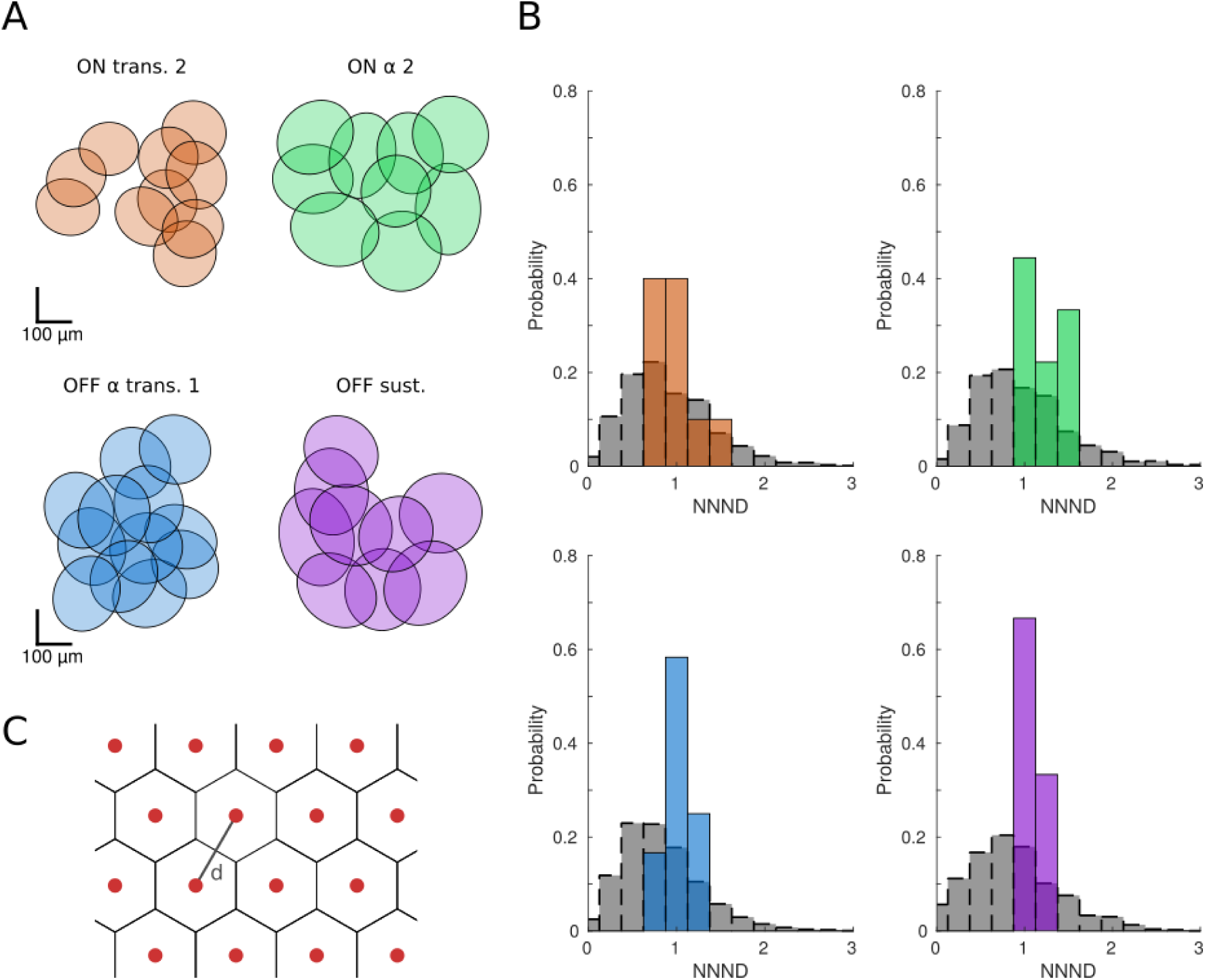
Mosaic Validation of the RGC types. **A**. Receptive field mosaics of four representative RGC types found by the chirp-based method. Ellipses represent the receptive field centers of the cells composing the mosaic. **B**. Normalized Nearest-Neighbor Distance (NNND) distributions for the mosaics shown in panel A. In each subpanel, we show the estimated distribution of nearest neighbor distances for the true mosaic (colored) and the same distribution for surrogate ensembles, generated by randomly sampling cells from different clusters (in gray). **C**. Tiling model used to estimate the recording bias of RGC types in MEA recordings. Ganglion cells of a same type (red dots) are assumed to be equally spaced at distance *d*, forming an hexagonal grid. Receptive fields of ganglion cells are modeled as adjacent, not-overlapping hexagons.

It is worth noting that this is a lower estimate of the number of well-isolated types that can be recorded with MEAs, since some clusters did not have enough cells to form good mosaics. This is expected since the MEA only covers an area of 0.25mm^2^. In this area, some types will appear only a few times. For example, if the cells of a given type have receptive fields of diameter ≈ 300*µm*, assuming little overlap between the different cells, there should not be more than 4 cells of that type in one recording. In addition, some ganglion cell types might be under or over-sampled by multi-electrode array recordings, due to the magnitude of their electrical activity or other physiological factors. To assess this, we made the assumption that ganglion cells are arranged in hexagonal grids with uniform spacing (fig. 2.C), and computed a recording bias Φ for each validated type. This index estimates whether a particular cell type is more (positive Φ) or less (negative Φ) likely to be recorded from with respect to other types (see Methods). Unsurprisingly, we found that all of the types for which a valid mosaic could be reconstructed in our recordings were over-sampled (table 1). In particular, ON *α*, ON low freq. and OFF sust. had a strong positive bias, meaning that these cells are more easily detected by MEAs in comparison to the other types.

To have a better insight of the biological types corresponding to our clusters, we compared our results to the classification carried out by [1] (fig. supplement 1, see Methods). For each validated cluster we identified the corresponding type in the [1] taxonomy based on the similarity of the responses to the chirp. When possible, we adopted the same nomenclature used by [1] (table 1). For some clusters it was not possible to establish a clear correspondence: in these cases, we just named our clusters according to the transience and polarity of their responses (ON trans. 1 and OFF sust.).

We then compared this “chirp-based” method with another clustering approach based solely on the RF (“RF-based”). We characterized each ganglion cell with the temporal profile and the diameter of its receptive field as previously, but not including the responses to the chirp this time. We applied the same procedure of dimensionality reduction and clustering mentioned above. The two methods compared here are in fact effectively equal in every aspect other than the choice of the physiological properties considered (see Methods). The RF-based method produced 15 clusters, and 6 of them could be validated as well-defined types, having at least one valid mosaic tiling uniformly the visual space.

We then compared the results obtained with our two approaches: chirp-based and RF-based. To make a quantified comparison we relied on the Matthews correlation coefficients (MCC), a measure commonly used to estimate how similar two clusters are, taking into account both false positives and false negatives. For both methods we looked at the mosaics of their well defined types, and used the MCC to find correspondences in the clusters generated by the other approach. We first considered the mosaics of the 6 types identified by the RF-based method: for each of them we could find a similar ensemble in the clusters of the chirp-based method (average MCC = 0.82 ± 0.13 std, worst match had MCC = 0.66). This indicates that all the types identified with the RF-based method can also be found by the chirp-based method. In particular, 5 (71%) of these chirp-based counterparts had also been validated as well-defined types, while the other 2 (29%) were discarded due to size or poor homogeneity in the responses to the chirp.

We followed the same procedure for the 15 types identified by the chirp-based method. In this case, it was not always possible to find well matching correspondences among the RF-based clusters (average MCC = 0.74 ± 0.16 std, worst match had MCC = 0.46, table 2). Of the 15 chirp-based mosaics, 5 (33%) were also identified and validated by the RF based approach (fig. 3.A). For the remaining 10 (66%), the RF-based best matching clusters seemed overpopulated and did not feature any mosaic organization (fig. 3.B). We also found 2 cases where ganglion cells from a single retina were pooled in an unique cluster by the RF-based method, while they were further split in two distinct valid types by the chirp-based method (fig. 3.B).

**Table 2:** Comparison of chirp-based and RF-based methods. For each type identified and validated with the chirp-based method, we found the corresponding type identified by the RF-based method. For each of these RF-based types, we report the size of the corresponding mosaic, whether it passed the mosaic validation test, and the similarity to the chirp-based counterpart (Matthew’s Correlation Coefficient). At the bottom: mean similarity across all the matches.

| RGC type | mosaic size | mosaic size | valid mosaic | similarity (MCC) |
| --- | --- | --- | --- | --- |
| ON trans. 1 | 8 | 15 | ✗ | 0.73 |
| ON trans. 2 | 10 | 9 | ✓ | 0.95 |
| ON trans. 3 | 13 | 11 | ✓ | 0.92 |
| ON low freq. | 10 | 9 | ✗ | 0.73 |
| ON DS sust. | 5 | 8 | ✗ | 0.79 |
| ON $\alpha$ 1 | 6 | 13 | ✗ | 0.68 |
| ON $\alpha$ 2 | 9 | 7 | ✓ | 0.88 |
| OFF step 1 | 7 | 20 | ✗ | 0.58 |
| OFF step 2 | 6 | 20 | ✗ | 0.54 |
| OFF $\alpha$ trans. 1 | 12 | 11 | ✓ | 0.96 |
| OFF $\alpha$ trans. 2 | 5 | 23 | ✗ | 0.46 |
| OFF slow | 7 | 8 | ✗ | 0.93 |
| OFF sust. | 9 | 14 | ✓ | 0.80 |
| OFF suppr. | 12 | 23 | ✗ | 0.63 |
| OFF $\alpha$ sust. | 5 | 9 | ✗ | 0.59 |

**mean similarity (MCC)** $0.741 \pm 0.16$

**Figure 3:**
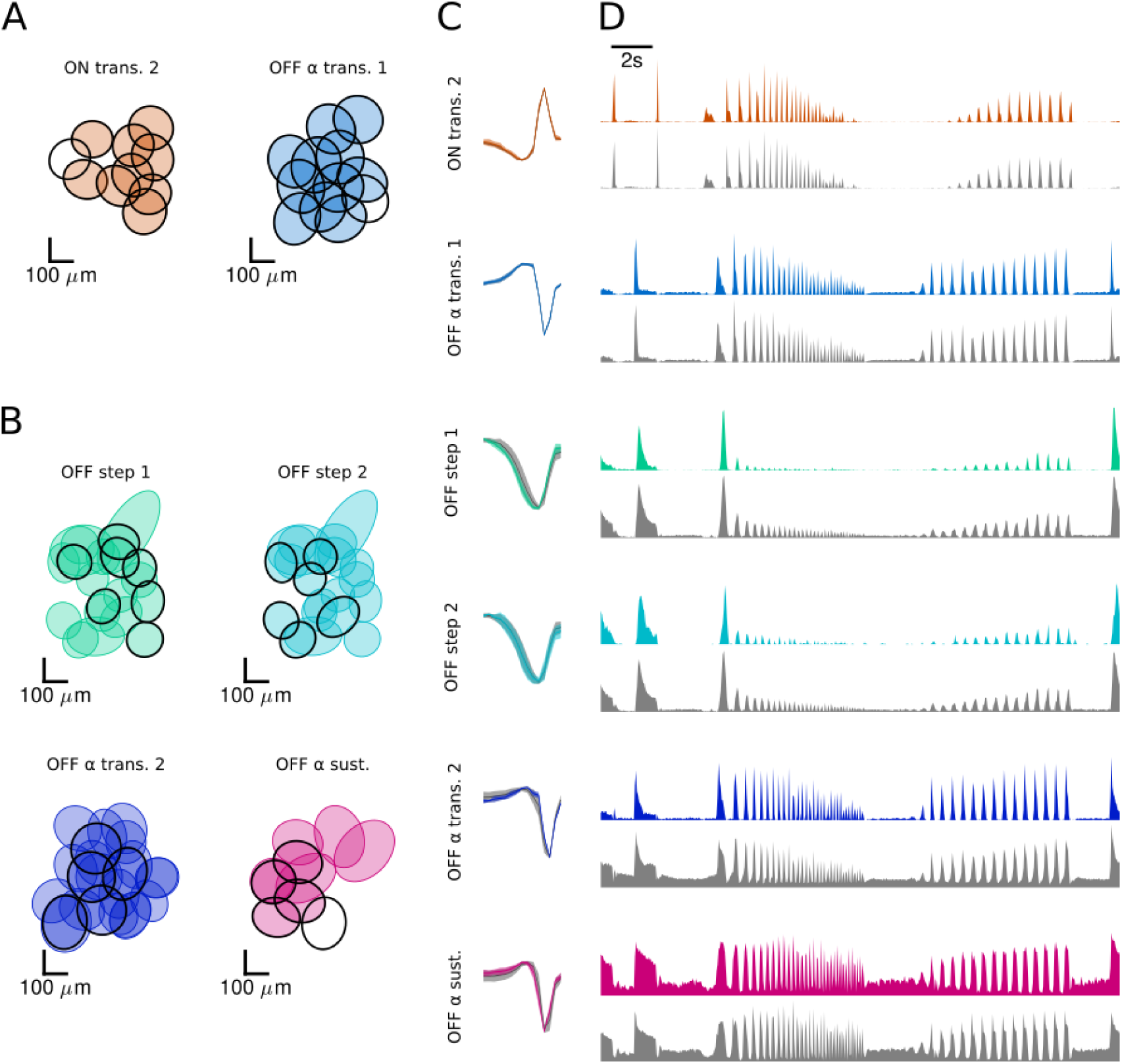
Comparison of chirp-based and RF-based methods. **A**. Mosaics of two representative RGC types found by the chirp-based method, superimposed with mosaics of the corresponding best matching types by the RF-based method. The chirp-based mosaics are indicated with black ellipses. The RF-based counterparts are depicted with filled colored ellipses. **B**. Same as A for the four types for which the RF-based method provided the poorest matches (MCC *<* 0.5). To note that the best match for OFF step 1 and OFF step 2 is in fact the same cluster. **C**. Mean temporal component of the receptive field across all cells composing the mosaics shown in A and B. In color: mean trace of the chirp-based mosaics; the shaded area shows its standard deviation. In gray: mean trace of the corresponding RF-based mosaics, with the shaded area showing its standard deviation. **D**. Mean responses to the chirp stimulus across all cells composing the mosaics shown in A and B. In color, mean responses of the chirp-based mosaics; in gray, mean responses of the corresponding RF-based mosaics.

We then looked at the temporal component of the receptive fields and at the chirp responses of the cells composing these mosaics. We found that, for all types, the mosaics identified by the two methods had very similar mean receptive fields (fig. 3.C). Conversely, the mean responses to the chirp were not always comparable (fig. 3.D). This shows that the chirp-based method can succeed in isolating ganglion cell types in cases where the RF-based method would fail and that, more generally, receptive fields alone are not sufficient to exhaustively characterize cell types.

### 2.2 Characterization of nonlinearities enhances discrimination of RGC types

Why can a method based on the responses to the chirp stimulus have more separating power than one based on the receptive field? Previous works have shown that the responses to full-field stimuli like the chirp can be well predicted by a linear-nonlinear model [2, 14, 16]. This model is made of two components: a linear filter, which corresponds to the receptive field, and a static nonlinearity, which converts the convolution of the stimulus with the filter into a firing rate prediction (fig. 4.A). We estimated the parameters of the LN model for each ganglion cell. For the linear filter, we took the receptive field estimated from the checkerboard stimulus, averaged over space to only keep the temporal dimension. We then estimated the non-linearity by relating the output of the filter to the firing rate (see Methods). We found that this LN model predicted well the responses to the full-field chirp for the large majority of the cells composing our mosaics (mean pearson coefficient *ρ* = 0.69 ± 0.21 std).

**Figure 4:**
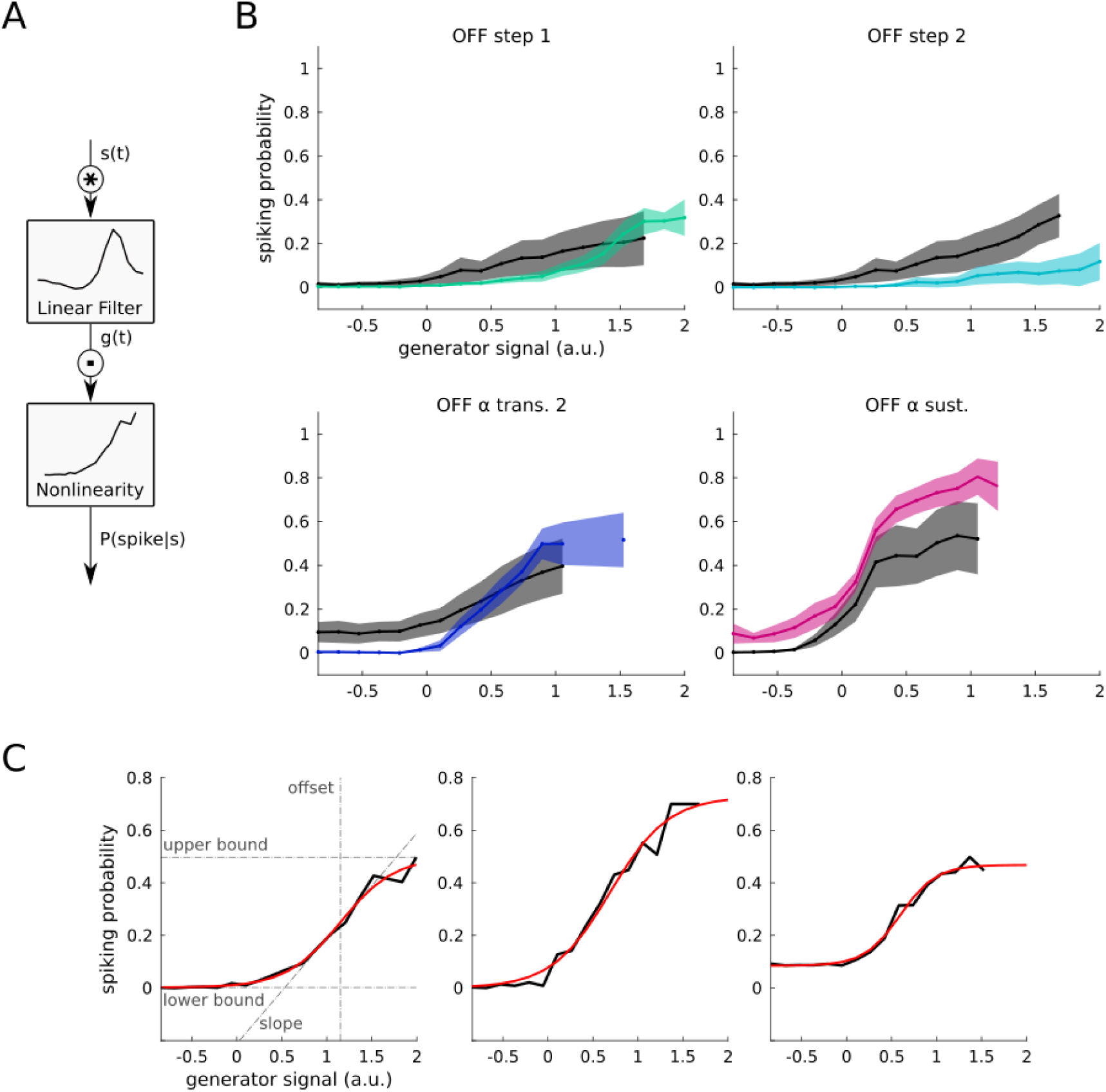
Comparison of RGC nonlinearities. **A**. The linear-nonlinear model used to characterize the nonlinearities of ganglion cells. The stimulus vector s(t) is convoluted with a linear filter, producing the generator signal g(t). This signal goes through a point-process non-linear function, which outputs the spiking probability P(spike s). **B**. Comparison between RGC nonlinearities found in the chirp-based and RF-based clusters respectively. Each subpanel shows the comparison for a different cluster (same ones as in fig. 3.B). In color, the nonlinearities of cells included in the cluster’s mosaic by both methods. In black, the nonlinearity of cells included only by the RF-based method. The bold line represents the average profile; the shaded area, its standard deviation. **C**. Three representative RGC nonlinearities fitted with the sigmoid function Θ. In black, the empirical non-linearity, computed using Bayes theorem; in red, its sigmoid fit. Dashed lines indicate the values of the four parameters of the sigmoid function: upper bound, lower bound, slope and offset.

Our results above show that, for some groups of cells, clustering solely based on the receptive field, i.e. the first component of the model, did not allow distinguishing different subtypes. Since the LN model is a compact descriptor of how ganglion cells will respond to the chirp, we thus hypothesized that these subtypes could be distinguished thanks to the other component of the model, i.e. the non-linearity. We focused on the 4 chirp-based types (OFF step 1, OFF step 2, OFF *α* trans. 1, OFF *α* sust.) for which the RF-based method provided the poorest matches (MCC *<* 0.6). Each of these chirp-based clusters was included in a RF-based cluster that also contained other cells. These RF-based ensembles were much larger than the chirp-based ones (average mosaic sizes respectively equal to 18.00 and 5.75) suggesting that they should have been further subdivided. We estimated the non-linearity for all the cells in these clusters. We then split each cluster obtained with the RF-based method in two groups: the cells who also belonged to the corresponding chirp-based cluster, and the cells who did not (fig. 4.B, see Methods). If our hypothesis holds true, the nonlinearities of these two subgroups should differ. To simplify their representation, we fitted the RGC nonlinearities with a sigmoid function with four parameters (fig. 4.C), and observed the distribution of these parameters in the two subgroups. For 3 types out of 4 (OFF step 1, OFF *α* trans. 1, OFF *α* sust.), these distributions were significantly different, meaning that the two subgroups indeed had distinct nonlinearities. This supports our hypothesis that the nonlinearity is a component that allows further division in different subtypes.

## 3 Discussion

We have compared two methods that have been used to separate ganglion cells in different cell types. The first one, based on their receptive field, allows to separate some types, but failed at separating others. In comparison, a method based on the responses to a standard ‘chirp’ stimulus could distinguish more cell types. To investigate the reasons for this better performance, we trained a LN model to predict the responses to the chirp stimulus. We found that some ganglion cells could have similar receptive fields, but different static nonlinearities, and as a consequence, distinguishable responses to the chirp. Methods that will take into account the non-linear responses of ganglion cells are thus more likely to distinguish different cell types.

Previous theoretical works have suggested that some ganglion cells with similar RFs may have different nonlinearities, and could thus correspond to different types [13]. However, direct evidence that nonlinearities could help dividing into further sub-types was lacking. Our study confirms this hypothesis and shows that ganglion cells can have similar receptive fields and different static nonlinearities, and therefore belong to different types.

Compared to the method of [1], here we only used the chirp to classify ganglion cells, while they used the responses to a larger ensemble of stimuli (flashes at different colors, moving bars, etc). Including responses to several distinct stimuli would certainly help further distinguishing different cell types. Here we restricted ourselves to the chirp stimulus because it was possible to relate the responses of the chirp stimulus to the receptive field easily: the only missing component was the static nonlinearity.

Characterization of non-linearity is thus necessary to define types on a functional basis. Including further characterization of the non-linear processing should help to do further division in cases where the chirp based stimulus was not able to distinguish different cell types. Two studies classified cell types based on their responses to a sequence of moving black and white stripes [3, 5]. However, there was no direct comparison with the method of [1]. A recent work by [10] did a full characterization using, among others, the responses to discs of various sizes. Since different cell types show a large diversity of suppressive surrounds [7], this additional characterization of nonlinearities in ganglion cell responses should probably help distinguish cell types.

## 4 Materials and Methods

### Multi-Electrode Array

MEA recordings were obtained from 6 *ex-vivo* isolated flat mounted retinae of wild type mice. Mice were sacrificed by quick cervical dislocation, and eyeballs were removed and placed in Ames medium (Sigma-Aldrich, St Louis, MO; A1420) bubbled with 95% O2 and 5% CO2 at room temperature. Isolated retinas were placed on a cellulose membrane and gently pressed against a MEA (MEA256 30 iR-ITO; Multi-Channel Systems, Reutlingen, Germany), with the RGCs facing the electrodes. Recordings were processed with Spyking Circus [24] to identify RGCs and sort their responses.

### Light Stimulation

In all the experiments we played a white noise stimulus (30Hz) to estimate the receptive fields of the ganglion cells. Each frame of this stimulus consists of a checkerboard of 38 by 51 checks, with each check of size 50*µ*m. The luminance of each check varies randomly at every frame according to a normal distribution.

To characterize the responses of our cells, we used a full field chirp stimulus (40Hz). This stimulus is composed of a first part in which a 1 second flash of white light is interleaved with 1 second of dim background light. The second part of the stimulus is composed of two ramps lasting 8s each: during the first one, luminance oscillates from dark to bright with constant contrast and increasing frequency; during the second one, it oscillates with constant frequency and increasing contrast (see fig. 1.A).

### Receptive Fields

To estimate the receptive field centers of the ganglion cells, we computed the spike-triggered averages (STA) of their responses to the white noise stimulus over an interval of 700ms (21 frames antecedent the spike). For each cell, this gives us a 3-dimensional matrix (2 spatial and 1 temporal components) representing the average check luminances of all the frames preceding a spike. We extracted the spatial component of the STA by computing its standard deviation of the mean over time. As a result, we obtained a 2-dimensional heat map that describes which checks evoke a response into the RGC. We fitted a 2-dimensional gaussian distribution onto this spatial component, and modelled the receptive field center as the ellipse delimited by one standard deviation of the distribution. Cells for which a good fit of the gaussian distribution could not be obtained were excluded from our dataset. To compute the temporal component of the receptive fields, we applied a spatial mask to the spike-triggered average considering only the checks lying inside the receptive field center, and averaged across space.

### Classification

We applied two clustering algorithms, one chirp-based and one RF-based, to the same dataset of 497 RGCs from 6 different experiments. Each clustering algorithm was composed by the three following steps:

#### [1] Feature extraction

For each retinal ganglion cell we computed the peri-stimulus time histogram (PSTH) of the responses to the chirp stimulus, normalized by dividing by its maximum (time bin = 50ms). We also computed the temporal component of the receptive field and the diameter of its center as described above. We then built the feature matrix of the cell population, which is composed of a feature vector for each ganglion cell. In the RF-based method, the feature vectors were obtained by concatenating the temporal RF vector and the diameter of the RF center. In the chirp-based method, we concatenated temporal RF, center diameter and PSTH to the chirp responses. We then reduced the dimensionality of the feature matrix through principal component analysis. We observed that the first 15 principal components were enough to describe more than 99% of the variance of the matrix. We hence generate a reduced feature matrix by pooling the coefficients of the 15 first principal components of each feature vector.

#### [2] Clustering

We then proceeded to the clustering of our data using a recursive unsupervised algorithm. We used expectation-maximization to fit a multivariate gaussian distribution to the data points constituted by the feature vectors of our cells. The number of gaussian components K was chosen maximizing the Akaike information criterion. At the end of this step, we assigned each cell to the gaussian component describing its feature vector best, obtaining K clusters of ganglion cells. We repeated this procedure recursively, subdividing each cluster into more subclusters until a termination condition was met. The termination conditions we considered are three:

**condition 1:** The cluster is too small to be further subdivided. This condition is met if the cluster size is smaller than the threshold *size*_*min*_.

**condition 2:** The cluster is consistent enough and it does not require to be further subdivided. Consistency of the cluster was assessed based on the homogeneity of the PSTHs and temporal RFs of its members. To quantify this consistency we built the matrices *PSTH*_*cluster*_ and *RF*_*cluster*_ by pooling respectively the PSTHs and temporal RFs of all the cells composing the cluster. We then computed the consistency index for both these matrices as follows:

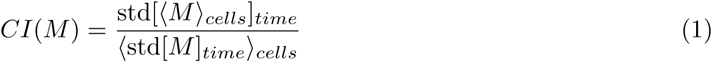

where M is the matrix considered (*PSTH*_*cluster*_ or *RF*_*cluster*_) of dimensions time steps by cells, and

⟨ ⟩ _*x*_ and std[ ]_*x*_ denote respectively the mean and standard deviation across the indicated dimension. For the chirp-based approach, the condition is met if:

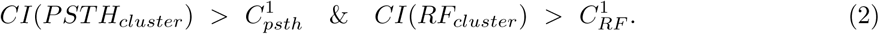

where 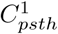 and 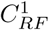 are two constants (see values below). For the RF-based approach, only the consistency of the temporal RFs was assessed, and hence the condition is met if:

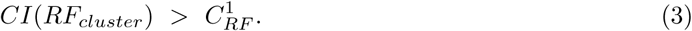

**condition 3:** Further subdivision of the cluster produces bad results. We subdivide the cluster and check the consistency of the subgroups. For the chirp-based method the condition is met if, for every subcluster, its size is smaller than *size*_*min*_ or if:

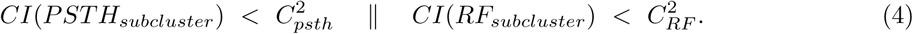

where 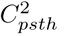 and 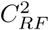 are two constants (see values below). For the RF-based method the condition is met if, for every subcluster, its size is smaller than *size*_*min*_ or if:

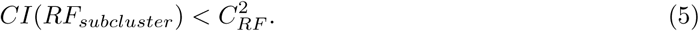

At the end of the process, all clusters with size smaller than *size*_*min*_, or *RF* consistency below 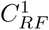 were discarded. For the chirp-based method, we also discarded all clusters with *PSTH* consistency below 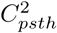. The results presented in this paper are obtained for *size*_*min*_ = 5, 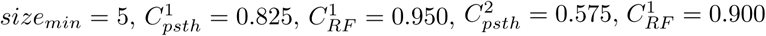. With the chirp-based method, we obtained a total number of 24 different cell clusters. With the RF-based method, we obtained 15 clusters.

#### [3] Validation

Retinal ganglion cells of the same type are spatially arranged in mosaics to maximize the coverage of the field of view and minimize intersections of their receptive field centers. We used this knowledge to validate our clusters. As mentioned above, our total ganglion cell population included cells from 6 different retinas. For each cell cluster then, we grouped together cells belonging to the same retina. For each of these groups, we superimposed the receptive field centers of its members, generating the corresponding mosaics (fig. 2.A).

We considered a cell cluster as valid if it featured at least one valid mosaic. Validity of mosaics was assessed as follows: a valid mosaic should have an uniform spacing of receptive fields of its composing elements. This means the distance between receptive fields centers of closest neighbours should be approximately the same for every member of the mosaic. For each mosaic member we calculated the normalized distance to its closest neighbour using the following formula:

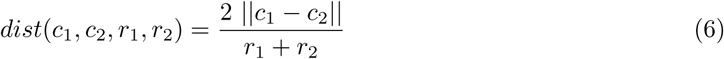

Where *c*_*i*_ and *r*_*i*_ represent respectively the center coordinates and the radius of the receptive field of a cell *i*.

We then built the histogram of normalized nearest-neighbour distances (NNNDs) to estimate the probability distribution of these distances for each mosaic. Good mosaic should feature unimodal distributions with a sharp peak around 2, entailing maximum coverage of the visual space and low intersection between receptive field centers. We used the Kolmogorov-Smirov (KS) statistical test to establish whether these distributions differed significantly from the ones computed on groups of cells sampled randomly. For each mosaic to validate, we generated 1000 surrogate mosaics of equal size by randomly sampling cells from the same retina population. We computed the same distance histograms for the surrogate mosaics, and averaged them to obtain an estimate of the NNNDs distribution for randomly sampled mosaics. We ran the one-sided Kolmogorov-Smirov test to check whether the cumulative NNNDs distributions in the real mosaics were smaller than the cumulative distribution in random mosaics. We considered as valid all the mosaics for which the KS test gave a value of *P <* .05. Following this procedure we validated 15 cell types identified with the chirp method (table 1), and 6 types for the RF-based method.

### Methods Comparison

To compare the results of a classification method *A* with respect to another method *B*, we used the Matthews Correlation Coefficient (MCC), an index that assesses the similarity of two sets in terms of precision and recall. First, we looked at all the types obtained with method *B*, which constitutes our ground truth against which we must compare the clusters from method *A*. We considered all the valid mosaics in these types, and looked for correspondences among the clusters found by method *A*. To find the best match of a mosaic 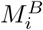 from method *B*, we looked at every cluster 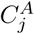 from method *A*, and defined the corresponding mosaic 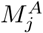 as the subset of 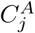 formed by only cells coming from the same retina as 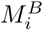 We then computed the MCC between 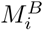 and all the mosaics 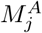, and selected the best match as the one for which MCC is maximized. For each RGC type *i* in *B*, the corresponding type in *A* is then found as:

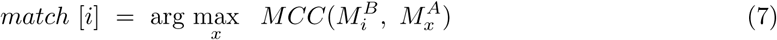

The overall performance of method *A* with respect to method B is then computed as the average MCC between all the mosaics in *B* and their best matches. An average MCC close to 1 indicates that most of the mosaics in *B* are accurately represented by clusters in *A*. Conversely, an average MCC close to 0 means that the mosaics in B are not well identified by *A*. The performance of the chirp-based method with respect to the RF-based method was 0.82, while the performance of the latter with respect to the former was 0.74. This means that the chirp-based method could identify the types validated by its competitor better than the RF-based method, indicating that the chirp-based method performs overall better (table 2).

### Correspondence to Biological Types

To have a better insight on which biological types corresponded to our clusters, we matched the clusters found by our methods to the RGC types identified by Baden and collaborators in [1]. These classifications were not directly comparable for two reasons: first, [1]’s classification was carried out on recordings of [*Ca*^2+^] activity (while ours was done on MEA recordings); second, the chirp stimulus used in the two studies was not identical. We then performed this comparison in three steps: first, we generated surrogates of [*Ca*^2+^] responses for each of our clusters. Second, we extracted snippets of [*Ca*^2+^] responses for both our and [1]’s clusters that were directly comparable. Third, we found correspondences between our clusters and the ones from [1] based on the similarity of the [*Ca*^2+^] traces (fig. supplement 1).

To compute the surrogate [*Ca*^2+^] responses we followed the same procedure described in [1]: for each validated cluster *i* we considered its cluster-mean response to the chirp 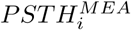 (fig. 1.A); we convoluted these traces with a filter *g* representing the average [*Ca*^2+^] event triggered by a single spike (see [1]), hence obtaining the surrogate [*Ca*^2+^] response 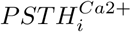:

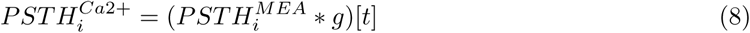

Then, we extracted for each cluster the 3 snippets of [*Ca*^2+^] responses that were directly comparable: an ON component (1*s* time interval after onset of the first flash), an OFF component (1*s* time interval after offset of the first flash), and a ramp component (22*s* time interval covering the two ramps). We then built the [*Ca*^2+^] vector 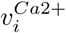 as the concatenation of these three snippets (fig. 5).

Finally, we used the Matthew Correlation Coefficient assess the similarity of the cluster vectors 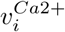 across the two classifications and find the type correspondences. We named each of our clusters after the most similar type in [1]’s taxonomy (fig. 1.C, table 1). When this correspondence could not be found unambiguously we named our types based solely on the polarity and transience of their responses (ON trans. 1 and OFF sust.). Note that following this method one of our clusters was assigned to the direction selective type ON DS sust. We named this cluster accordingly, although the direction selectivity of its members has not been verified directly.

### Multi-electrode Array Recording Bias

To establish whether multi-elecrode array recordings are biased towards some particular ganglion cell types, we estimated a recording bias for all the RGC types found with the chirp method. We defined the bias Φ_*i*_, for a cell type *i* as:

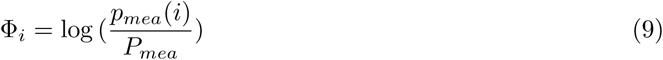

where *p*_*mea*_(*i*) is the probability of being recorded for a cell of type *i*, and *P*_*mea*_ is the probability of being recorded for a generic ganglion cell, regardless of its type. This index estimates whether cells belonging to a particular type are more (or less) likely to be detected by multi-electrode arrays with respect to the others: a positive bias entails that the type is over-sampled, while a negative bias means the type is under-sampled. We computed the probabilities *P*_*mea*_ and *p*_*mea*_(*i*) respectively as:

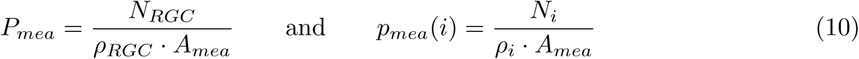

where *N*_*RGC*_ is the total number of RGCs recorded (in a given experiment), *N*_*i*_ is the number of RGCs of type *i* recorded, *ρ*_*RGC*_ is the average planar density of retinal ganglion cells, *ρ*_*i*_ is the estimated planar density of cells of type *i*, and *A*_*mea*_ is the area of the multi-electrode array. Note that the value of *A*_*mea*_ is not relevant to the computation of the bias, as it cancels out in equation 9. For *ρ*_*RGC*_, we chose a value of *ρ*_*RGC*_ = 5000*mm*^*−*2^, consistent with what is reported in the literature [4]. To estimate *ρ*_*i*_, we assumed that all cell types tile the visual field forming hexagonal grids (fig. 2.C). Following this model, the density of an RGC type can be computed as the inverse of the area of its hexagonal tiles:

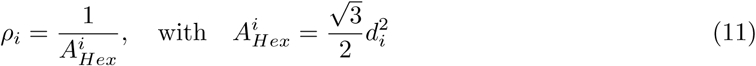

where *d*_*i*_ is the average distance between two neighbouring cells of type *i*. We computed *d*_*i*_ as the median nearest neighbour distance across cells of type *i*.

### Modeling Of Nonlinearities

To analyze the non-linear component of the RGC responses to the chirp stimulus we used a linear-nonlinear model. This model is composed of a linear filter which convolutes the stimulus, producing a generator signal *g*, and a point-process nonlinearity that turns the generator signal into the spiking probability *p*(*spike* | *g*) (fig. 4.A). As the chirp stimulus is full-field, it can be just represented with a one-dimensional vector of luminance over time. As a result, also the linear filter is one-dimensional, and it can be well approximated by the temporal component of the RF. To estimate the empirical nonlinearity of the model *p*(*spike*|*g*) we made use of the Bayes theorem:

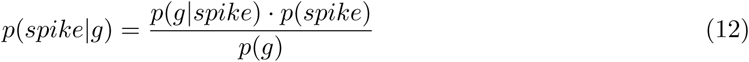

To estimate the prior probability *p*(*g*), we convoluted the linear filter with the chirp stimulus vector (normalized between zero and one) to obtain a snippet of generator signal. We calculated *p*(*g*) as the histogram built on this generator signal. To estimate the marginal probability *p*(*spike*), we looked at the responses of the cell to each repetition of the chirp stimulus. We computed this probability as the number of chirp frames producing (at least) a spike, divided by the total number of frames. To estimate the posterior probability *p*(*g* | *spike*), we computed the generator signal for the chirp stimulus as above. Then, we looked at the responses of the cell to each repetition of the chirp stimulus. For each chirp frame for which at least one spike was produced, we stored the corresponding value of the generator signal. We then calculated *p*(*g* | *spike*) as the histogram built on these generator signal values. To get a simpler representation of the spiking probabilities *p*(*g* | *spike*), we fit the empirical nonlinearities obtained above with the sigmoid function Θ shown below:

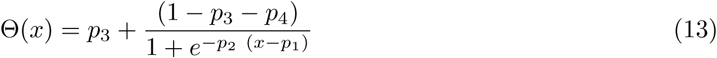

Where *p*_1_, *p*_2_, *p*_3_, *p*_4_ represent respectively the offset, the slope, the lower bound and the upper bound of the curve (fig. 4.C).

### Comparison Of Non-Linearities

To test the hypothesis that the RF-based method fails at identifying RGC types due to its inability to discriminate cell nonlinearities, we considered the four RGC types (OFF step 1, OFF step 2, OFF *α* trans. 1, OFF *α* sust.) for which the comparison between RF-based and chirp-based methos gave the poorest results (MCC *<* 0.6). Given the pair 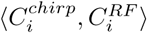, representing respectively a cluster obtained with the chirp-based method and its best match obtained with the RF method, we built the set of cells belonging to both clusters *S*^*both*^ and the set of cells belonging only to the RF-based match *S*^*RF*^. For each of these cells, we considered the parameter vector ***p*** composed by the four parameters of the corresponding sigmoid fit. We used multivariate analysis of variance (MANOVA) to estimate the multivariate distribution of the parameter vector ***p*** in the two sets *S*^*both*^ and *S*^*RF*^, and test whether these distributions were significantly different. We found that for 3 out of 4 types the difference was significant (OFF step 1: *P <* .001, OFF *α* trans. 1: *P* = .041 and OFF *α* sust.: *P* = .028 respectively), while for the 4th type it was not possible to draw any conclusion (OFF step 2: *P* = .085).

---

**Figure supplement 1:**
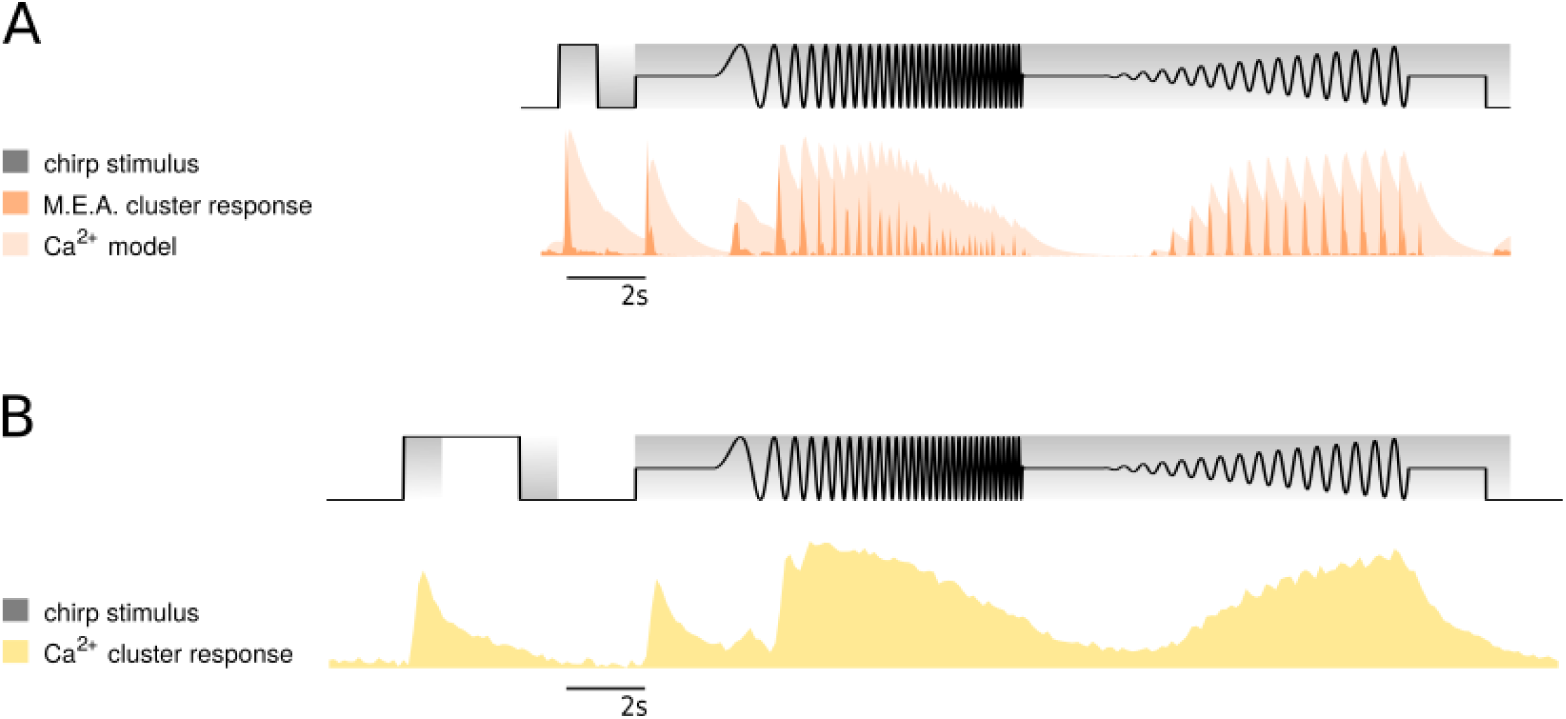
Comparison with Baden et al. RGC classification. **A**. On top: the temporal profile of our chirp stimulus. Below, in dark orange: the normalized mean response (firing rate) of a representative RGC type (ON trans. 3). In light orange: model of the calcium trace for the same response. **B**. On top: the temporal profile of the chirp stimulus used by [1]. Shaded regions correspond to the temporal snippets considered for type matching. Below, mean calcium trace of the corresponding type (ON transient) identified by [1].

## Notes

### Competing Interest Statement

The authors have declared no competing interest.

